# Statistics used to infer inter-breeding between humans and Neanderthals are strongly predicted by flanking sequence heterozygosity

**DOI:** 10.1101/2020.10.11.334862

**Authors:** William Amos

## Abstract

A large and rapidly expanding literature has grown out of the observation that humans carry a genetic legacy reflecting ancient inter-breeding with archaic hominins such as Neanderthals and Denisovans. However, a recent study suggests that a commonly used statistic used to assess legacy size, D, is driven mainly by heterozygous sites in Africa acting to increase divergence from our common ancestor rather than introgressed fragments outside Africa reducing divergence. To test this new model, I analysed how D is influenced by heterozygosity within a kilobase of each putative introgressed base. I find that flanking heterozygosity is a potent predictor of D, with introgression always being inferred as having occurred into the population with lower heterozygosity. This pattern cannot be driven by any introgressed fragments themselves, which simulations show create the exact converse pattern, but instead appears to be generated by heterozygosity acting to drive increased divergence from the ancestral sequence. This new model explains why introgression of haploid or semi-haploid regions is essentially lacking and why introgression is often inferred around immune genes and other regions under strong selection. More generally, these results raise the possibility that reported legacies are largely an artefact arising out of the false assumption that mutation rate is constant.

## Introduction

The idea that humans inter-bred with related lineages such as Neanderthals and Denisovans (Green *et al*. 2010; Qin and Stoneking 2015; Dannemann *et al*. 2016; Sankararaman *et al*. 2016; Vernot *et al*. 2016) was accepted rapidly and has become the focus numerous studies and high profile publications. The discovery of hybrid skeletons (Fu *et al*. 2015; Slon *et al*. 2018) lends direct support to the idea that inter-breeding did occur and yielded viable offspring. Analyses of aligned genomes has allowed the size of the resulting genetic legacies to be inferred (Sankararaman *et al*. 2014), with Eurasian genomes estimated to carry around 2% Neanderthal DNA (Mallick *et al*. 2016; Sankararaman *et al*. 2016), a value that rises from west to east (Wall *et al*. 2013), and even higher levels of Denisovan DNA found in some Oceanians and Australians (Meyer *et al*. 2012; Vernot *et al*. 2016; Jacobs *et al*. 2019).

Despite the seemingly overwhelming evidence in favour of ubiquitous archaic legacies in non-Africans, some aspects remain unexplained. As yet, there have been no reports of archaic mitochondrial DNA or Y chromosomes in humans and any legacy on the X chromosome is minimal (Sankararaman *et al*. 2014; Sankararaman *et al*. 2016). Such absences might reflect selection (Green *et al*. 2010), though it is not obvious how selection could be both strong enough to prevent these genomic regions introgressing, yet at the same time weak enough to allow hybrid individuals to survive, prosper and leave many descendants. More directly, a recent study shows that the signal widely assumed to reflect introgression resides largely with heterozygous bases in Africans rather than the pattern one would expect from introgression, where the signal should rest with heterozygous bases in *non-Africans* (Amos 2020). This pattern is best explained by a model in which faster evolution in Africans causes increased divergence from our common ancestor rather than introgressed DNA outside Africa acting to reduce divergence.

A higher mutation rate in Africans is unexpected, but evidence is mounting in support of a model in which mutation rate is elevated at and around heterozygous sites (Rubinsztein *et al*. 1995; Tian *et al*. 2008; Amos 2010; Yang *et al*. 2015) such that the large loss of heterozygosity associated with the out of Africa bottleneck (Prugnolle *et al*. 2005) caused a parallel drop in mutation rate. Thus, microsatellite mutation rate is predicted by human population size (Amos 2011) and mutation rate is elevated at loci with higher heterozygosity even after controlling for allele number (Amos 2016). Equally, human populations differ in mutation rate (Mallick *et al*. 2016) and, when only variants that likely post-date the out of Africa event are considered, the mutation rate in Africa is appreciably higher than elsewhere (Amos 2013). Also, across the genome, the difference in mutation rate between Africans and non-Africans is strongly predicted by the amount of diversity lost (Amos 2013): relatively, the more heterozygosity was lost, the greater the excess mutation rate in Africa. Finally, there are highly significant differences between human populations in which specific mutational changes are more likely (Harris 2015; Harris and Pritchard 2017) and these differences are predicted by flanking sequence heterozygosity (Amos 2019), increasing the plausibility of a parallel link to mutation rate.

As a direct test of the idea that heterozygosity acts as an important driver of signals interpreted as evidence of introgression, I asked whether the distribution of putative introgressed bases could be predicted by heterozygosity in the immediately flanking region.

## Results

Genetic legacies were originally estimated using the so-called ABBA-BABA test (Green *et al*. 2010; Durand *et al*. 2011; Martin *et al*. 2015), applied to a four-way alignment of African – European – Neanderthal – chimpanzee, in that order. Bases are sought at diallelic sites where the Neanderthal and chimpanzee bases differ as do the two human bases. The chimpanzee base is always labelled A, giving two forms, ABBA and BABA, depending on which human matches the Neanderthal. The standardised difference in counts, calculated as (ABBA - BABA)/(ABBA + BABA) is referred to as D and can vary between -1 and 1. There is no convention concerning population order but D is usually calculated such that positive values indicate excess base-sharing between Neanderthals and Europeans compared with Neanderthals and Africans. In the absence of introgression and if mutation rate is constant, the expectation of D = 0.

### a) Impact of window placement

For a highly localised view on the possible impact of heterozygosity on D, I searched the 1000 genomes project Phase 3 data (1000 Genomes Project Consortium 2010) exhaustively for informative bases, i.e. diallelic loci where the chimpanzee and archaic genomes differ and hence that contribute to the numbers of ABBAs or BABAs. At each such base I calculated heterozygosity in both human population one and human population two, based of two symmetrically placed 1Kb windows either side of but excluding the focal base. The total 2Kb window size was chosen to reflect the fact that heterozygote instability is thought to operate via gene conversion-like events of the order of 1-2Kb in length. Three different window placements displacements from the focal base of 0Kb, 2Kb and 4Kb. ABBAs and BABAs were then counted according to the heterozygosity bins in which each base falls, yielding a 2D matrix of D values. To illustrate how D varies in these windows I chose the most extreme case of D(*Africa, East Asia, Neanderthal, chimpanzee*), the actual human populations illustrated being ESN (Africa) and CHS (East Asia), though other combinations yield essentially identical results.

Figure 1 depicts how D varies according to flanking sequence heterozygosity in Africa and East Asia, represented by ESN and CHB respectively. In the lefthand panel, heterozygosity is calculated over a 2Kb window centred on but excluding the informative base. The middle panel repeats similar data but here heterozygosity is calculated using two 1Kb windows placed at 4-5Kb either side. The two plots are very similar and reveal a pattern where D values above the diagonal, where heterozygosity is greater in East Asia, are largely negative (red), suggesting introgression into Africa, while D values below the diagonal D are largely positive, suggesting introgression into East Asia. The overall positive value reported in the literature (Green *et al*. 2010; Meyer *et al*. 2012; Sankararaman *et al*. 2014) seems to reflect the larger proportion of data points that fall below the diagonal, due to generally higher heterozygosity in Africa, rather than to a uniform increase in D across the entire plot. The righthand panel is a difference plot based on adjacent – displaced D values for all cells where both plots contain data. With purple representing negative differences and green representing positive differences, although weak, essentially the same pattern is produced. The intermediate window gives intermediate results, being weaker than the adjacent window but stronger than the 4Kb displacement. The meaning of this tendency for closer windows to produce stronger trends is discussed more below.

**Figure 1.**
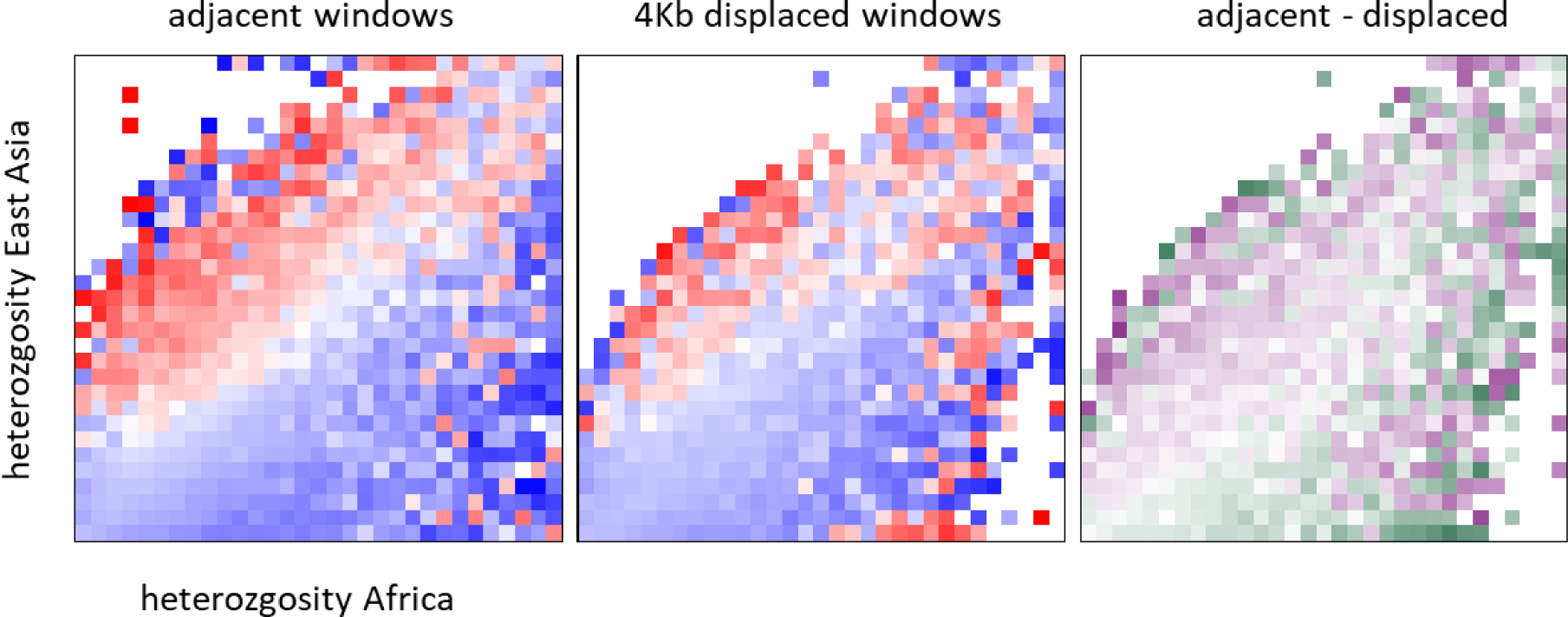
Dependence of D on heterozygosity: the impact of window proximity. A representative African – non-African population comparisons was selected (ESN – CHB) and all bases contributing to D(ESN, CHB, Neanderthal, chimpanzee) identified. Flanking sequence heterozygosity was estimated in 30 bins in both human populations for two 1Kb windows either side of each base, the windows being placed either adjacent or displaced by 4Kb. Each plot shows D, colour-coded according to value (from darkest red = -1 through white = 0 to darkest blue = 1), calculated for each combination of heterozygosity values where counts of ABBA+BABA exceed two. The righthand panel is a difference plot, calculated as D_adjacent_ – D_displaced_, with purple = negative and green = positive. Since most differences are small, the pattern is accentuated by taking the signed root of the absolute difference.

### b) Consistency across comparison types

The above process was repeated for SNPs and INDELs separately and also for both Neanderthals and Denisovans. Representative plots are presented in Figure 2 based on adjacent 1Kb windows (hereafter the default). The left-hand panel is based on SNPs for the Neanderthal comparison and therefore repeats the lefthand panel from Figure 1. The middle panel is the equivalent plot for Denisovans. The two different archaic plots are very similar, though the Denisovan plot tends to show a slightly more intense pattern, with larger magnitude D for any given displacement from the diagonal. Although subtle, this tendency is replicated across other population combinations. The third panel represents a plot based on heterozygosity calculated using INDELS alone, with Denisovans chosen over Neanderthals because they yield a somewhat stronger pattern. INDEL plots understandably carry a much narrower range of heterozygosity values compared with SNPs, even after multiplying their scores by four (as done in this figure) but generally show a pattern that is consistent with equivalent plots based on SNPs. Most INDEL plots appear to show a weaker trend for increasing magnitude of D away from the diagonal (the example shown is somewhat stronger than average) but it should be remembered that to make the plot directly comparable to SNPs, the multiplication would be removed, making the spread of points 4-fold narrower.

**Figure 2.**
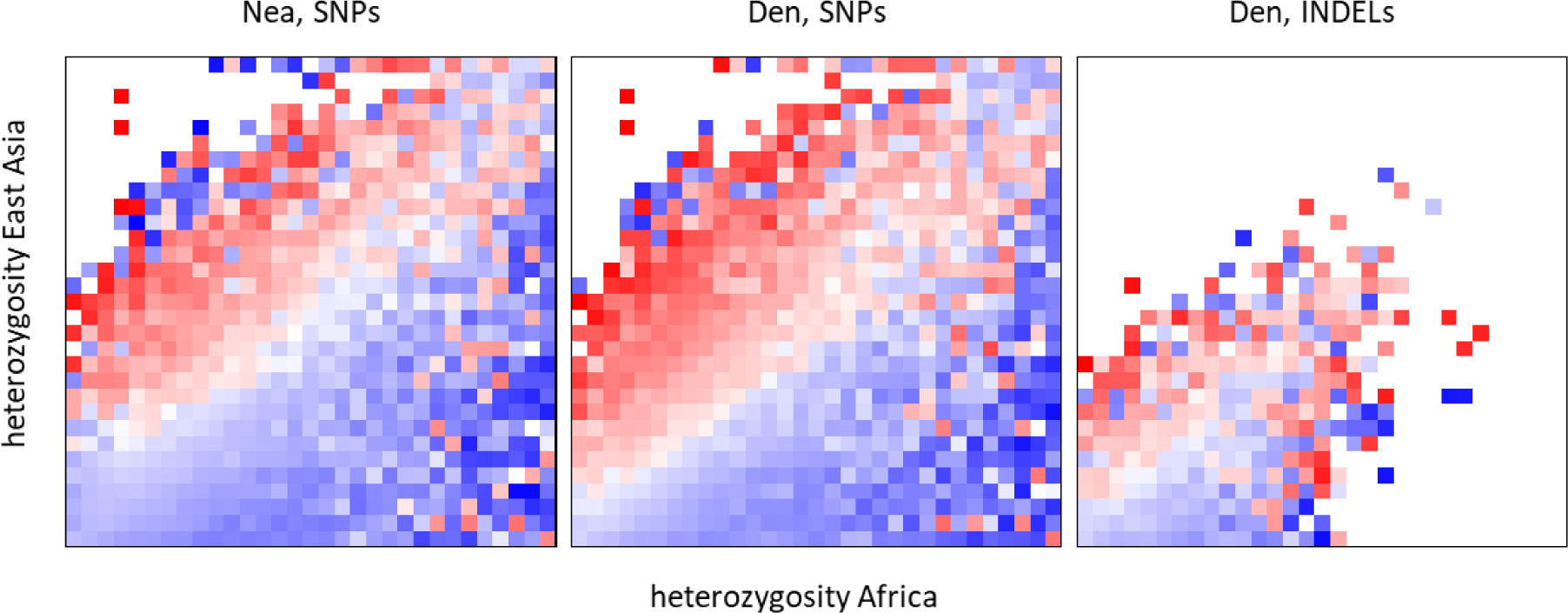
Dependence of D on archaic species and variant type used to calculate heterozygosity. The three plots are generated as described in the legend to Figure 1. The lefthand panel is the same and is included for reference. The middle panel is the same but with the Denisovan instead of the Neanderthal. The righthand panel is the same as the middle panel except that instead of SNPs, heterozygosity is estimated using only variants classified as indels.

### c) Comparisons between different population combinations

To explore the consistency of these patterns across the various pairwise population comparisons, I averaged D across all possible comparisons within each combination of the five major geographic regions: Europe (EUR), East Asia (EAS), South Asia (SAS), Africa (AFR) and America (AMR). Results are summarised in Figure 3, with data for Neanderthals above the diagonal and for Denisovans below the diagonal. Within-region comparisons are only presented for Neanderthals but the equivalent Denisovan plots are near-identical. As expected, comparisons within a region exhibit much narrower profiles, reflecting the much smaller range of heterozygosity differences present. As seen before, the Neanderthal and equivalent Denisovan profiles are all extremely similar. In all cases, the core pattern is largely preserved, with D values above the diagonal becoming increasingly negative (red) with increasing distance from the diagonal and points below the diagonal becoming increasing positive (blue). However, the pattern is far from perfect. Some of the largest absolute values (strongest colours) appear far from the diagonal on the ‘wrong’ side, a good example being the strong red that appears in the bottom right of African – American comparisons. This is particularly noticeable in general for comparisons involving American populations, including comparisons within America. Finally, notice how the ordering of the populations has relatively little impact. For EUR, EAS and SAS, D is calculated as D(region, AFR, NEA, CMP) with Africa on the X-axis, while for America, Africa is on the Y-axis. The core pattern remains similar even though for AFR-AMR the higher African heterozygosity causes data to spread upwards towards the top left rather than rightwards towards bottom right.

**Figure 3.**
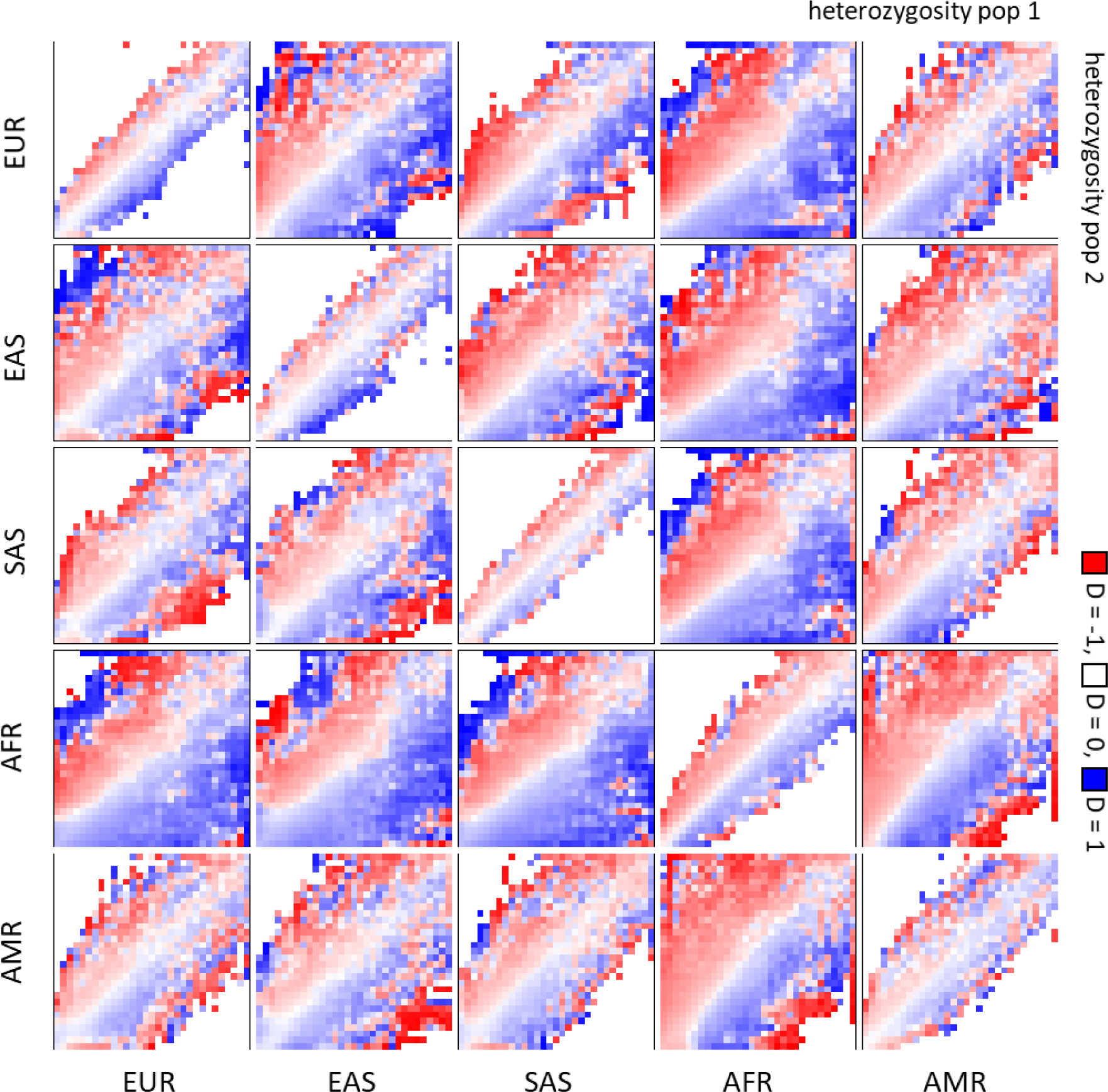
Relationship between D and flanking sequence heterozygosity for regional population comparisons. All combinations of populations were analysed using 1Kb heterozygosity windows adjacent to each informative base. Data are then averaged across all unique population comparisons from the same regions (e.g. all European populations against all East Asian populations). D(*p1, p2, Neanderthal, chimpanzee*) is calculated for each heterozygosity bin combination, *p1* being the X-axis population and *p2* being the Y-axis population. Data for Neanderthals are presented above and including the diagonal. Data for Denosivans are presented below the diagonal. Within-region comparisons for Denisovans are not presented but show negligible differences from the equivalent Neanderthal plot.

### d) Simulations

In order to help interpret the empirical data, I ran a series of coalescent simulations using ms. Each simulation generates sequence data for a chimpanzee, a Neanderthal, two ‘African’ populations and two non-African (‘European’) populations. The root European population experienced a bottleneck and may or may not receive a 3.5% injection of Neanderthal DNA before splitting. In addition to varying whether or not introgression occurred I also varied the timing of the human – Neanderthal split (HNS) from 300,000 to 1 million years ago. This range reflects the rather recent value used by Green et al. amongst others as well as a older estimates based on dental evolution (GÓmez-Robles 2019) and a variety of genetic data (Stringer 2016; Hajdinjak *et al*. 2018; GÓmez-Robles 2019). Sequence data were generated as 5,000 x 500Kb blocks with recombination and analysed as for the real data.

Without introgression (Figure 4), plots for the simulated data resemble those for the empirical data. There is a general tendency for D values below the diagonal to be blue (positive, suggesting introgression into ‘Europe’) but this pattern weakens as the HNS is pushed back towards a million years. This trend makes intuitive sense. In simulated data based on an infinite sites model without introgression, all informative sites reflect incomplete lineage sorting. With a recent split time, alleles shared with Neanderthals will tend to be common and to reflect the ancestral state, such that homozygous regions will show enhanced affinity to this and hence to Neanderthals. Pushing back the HNS time allows more time for drift and so the derived allele frequencies increasingly approach a uniform random distribution (all frequencies equally likely), weakening or even removing the link between zygosity and ancestral state. As expected, the breadth of scatter decreases from left to right across the three panels, from the most diverged, Africa – Europe comparison, to the comparison between two related, bottlenecked populations in Europe.

**Figure 4.**
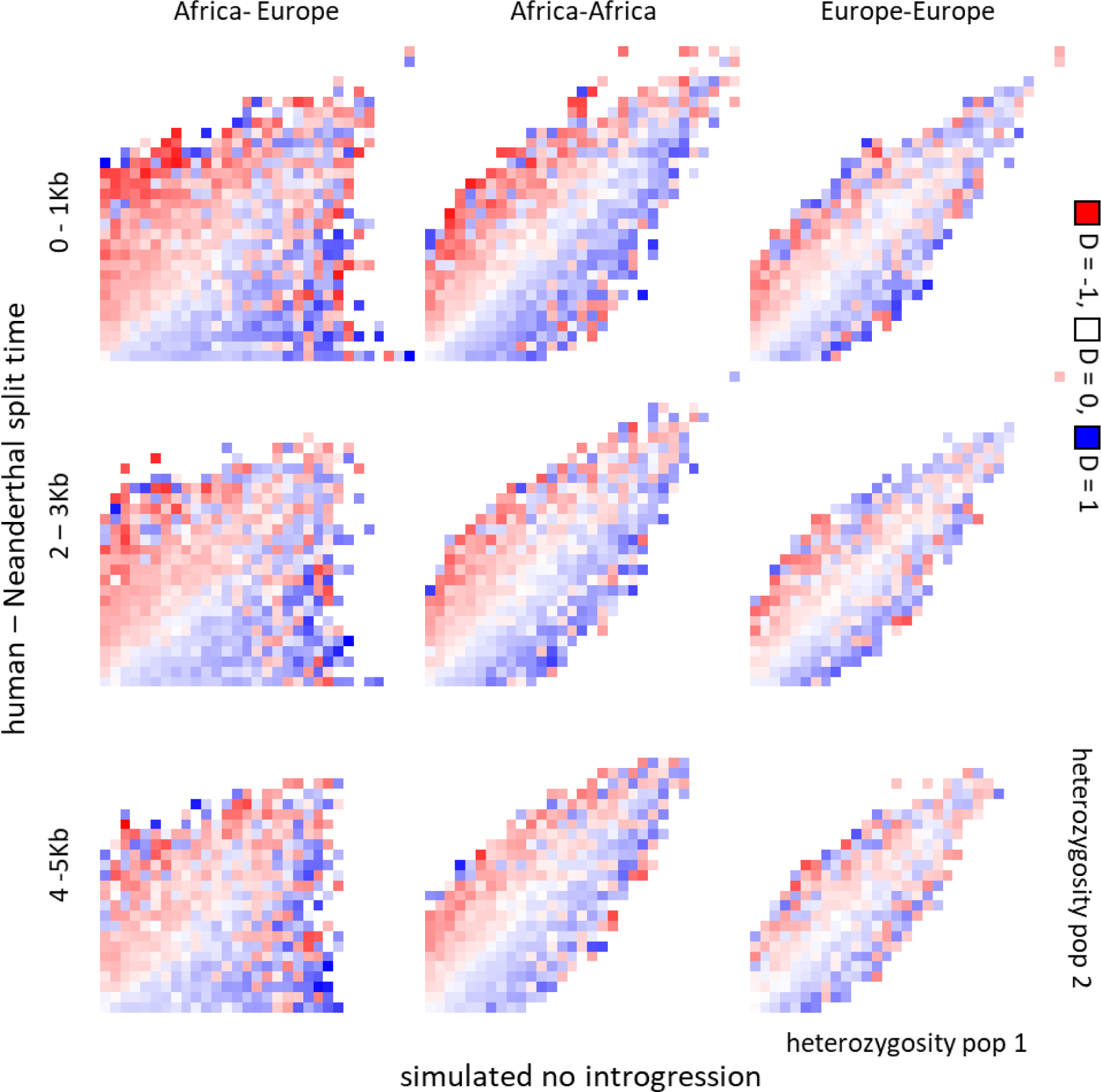
Simulated data without introgression. Data were simulated based on two ‘African’ and two ‘European’ populations, allowing both within and between region comparisons. As with the empirical analysis, three different heterozygosity window placings were analysed: adjacent and displaced by two or four kilobases.

Adding in introgression elicits a striking change (Figure 5). In line with theory (Durand *et al*. 2011), D values tend to increase in magnitude as the HNS is pushed back in time, due to the resultant reduction in numbers of background ABBAs and BABAs generated by incomplete lineage sorting. As expected, within Africa, introgression has no impact because I have not allowed migration to link Africa and Europe. Within Europe, the pattern is now reversed across the diagonal, with values below the diagonal now appearing red (negative) and above the diagonal being blue (positive). This agrees with expectations. Introgression around 50-60,000 years ago leaves little time for introgressed fragments to drift to high frequency. Consequently, most introgressed fragments should be found in the heterozygous state and D captures this trend: at sites where one population has higher heterozygosity than the other, D indicates that the population with higher heterozygosity has received more Neanderthal DNA. Lastly, in Africa – Europe comparisons the picture tends more and more towards being uniformly blue as the HNS is pushed back. Here, the impact of heterozygosity is rather swamped by two stronger trends: Africans carrying generally greater diversity and all introgressed fragments being found in Europe. Consequently, most D values are positive, indicating introgression into Europe. The reversal across the diagonal within Europe and tendency towards all values being positive in Africa-Europe comparisons are two trends that, while they make intuitive sense, appear completely absent from the empirical data.

**Figure 5.**
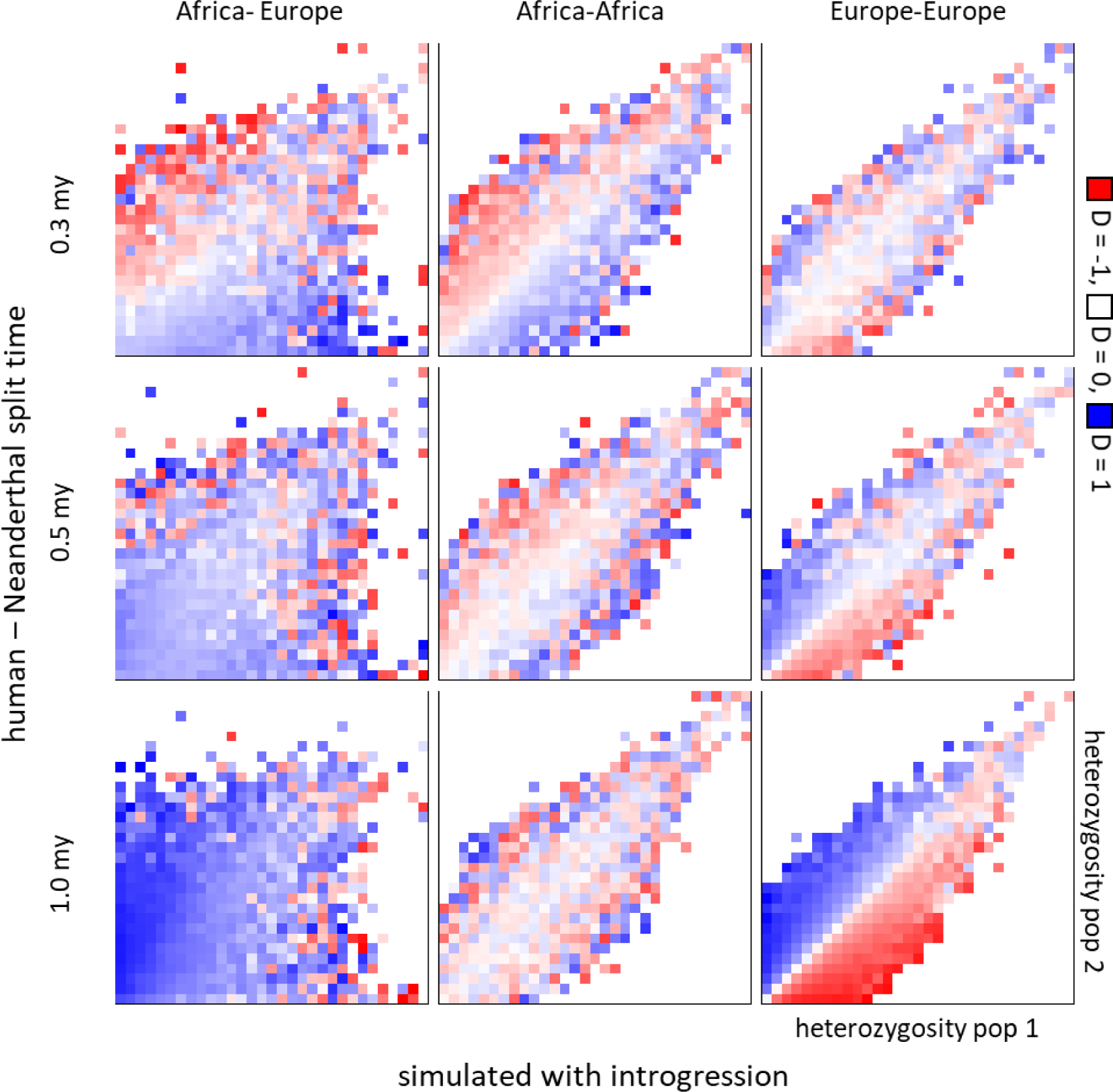
Simulated data with introgression. Data were simulated as in Figure 4 but with Neanderthal introgression into the basal European population. Also, instead of analysing three different window placements, here all analyses are conducted using the adjacent window for estimating heterozygosity. Three different human-Neanderthal split times were simulated: 0.3 mya, 0.5 mya and 1 mya.

Finally, I asked about the extent to which a link between D and heterozygosity declines as heterozygosity is estimated using more distant sites. Results are presented as difference plots between the adjacent and 4Kb displaced windows and, to stand best chance of seeing an effect, I used the oldest HNS time of one million years. In simulated data (Figure 6, top row), the differences are either negligible (within Africa, purple and green approximately randomly distributed) or tend to show generally higher values in the adjacent window (the two other panels, most values are positive, purple). Crucially, the reflected pattern across the diagonal is lost. In contrast, in all equivalent plots for the real data, the reflected pattern across the diagonal is preserved. As with the single example presented in Figure 1 above, wherever D is positive based on the adjacent window it tends to be smaller in the displaced window. Equally, wherever D is negative in the adjacent window it tends to be less negative in the displaced window. This confirms that the relationship between heterozygosity and D is noticeably stronger closer to the focal base.

**Figure 6.**
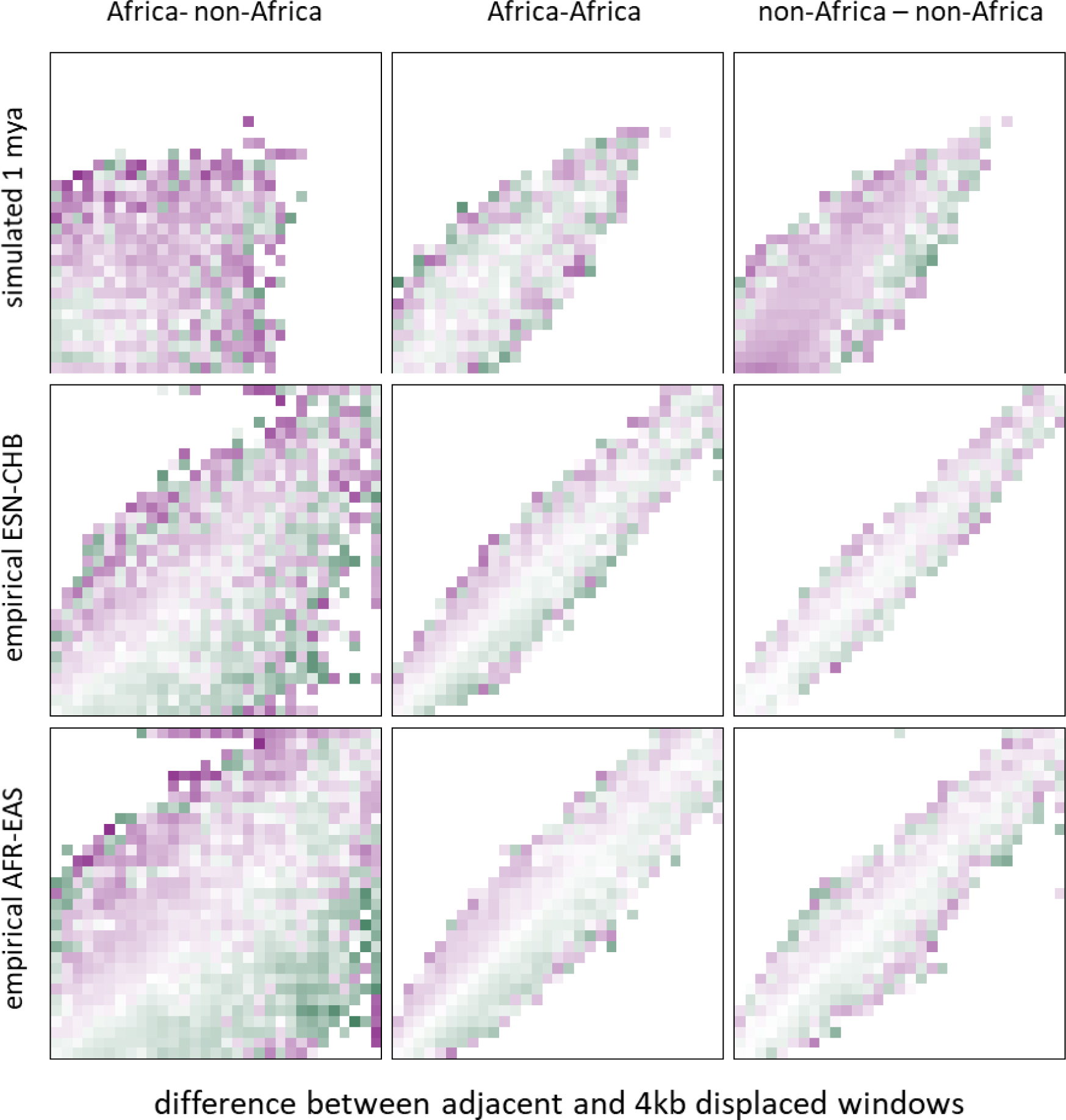
Effect of window placement. Each panel represents a difference plot for the adjacent – 4 kb displaced windows used for estimating heterozygosity. The top row depicts results for simulated data for the most extreme case of introgression with the most ancient human – Neanderthal split time of 1 mya. The middle row depicts empirical data for one particular pairwise population comparison, ESN - CHB. The bottom row is based on the average of all possible African – East Asian comparisons. Columns contain the three main comparisons: Africa – non-African, within Africa and non-African – non-African. Other details of difference plots are found in the legend to Figure 1

## Discussion

Here I report a widespread association between D, a statistic used to infer introgression, and local heterozygosity, as measured in the genomic region immediately surrounding each informative site. Similar patterns are seen for both Neanderthals and Denisovans and for heterozygosity measured using both indels and SNPs. The association increases in strength with proximity to the informative site. Importantly, despite some scatter, regardless of the population comparisons used, D always tends towards indicating archaic introgression into the population with *lower* heterozygosity.

There are at least four reasons for believing that an association between heterozygosity and D is not driven by any introgressed fragments themselves. Most obviously, the relationship goes in the exact opposite direction to the one expected, something that is confirmed by simulations. Introgressed fragments should on average be less similar to human sequences than are other human sequences from the same population. Consequently, where introgressed fragments are present, heterozygosity should tend to be higher in the population that carries them. In practice, where D is non-zero, it is the human where putative introgressed bases sit in regions with *lower* heterozygosity that appears closer to the archaic. Second, if introgressed fragments are the main driver, the relationship should be absent or at least much diluted inside Africa, yet the pattern remains strong. Third, introgressed fragments are expected to be of the order of 50Kb long (Seguin-Orlando *et al*. 2014), which is large relative to the 1Kb - 2Kb scale over which effects are seen. With fragments of this size, the difference between a window adjacent to an informative base compared with a window 2Kb away should be negligible, yet a clear difference exists. Finally, heterozygosity is an extremely dynamic property that will have changed much since any introgression occurred (Prugnolle *et al*. 2005). Consequently, unless introgressed fragments themselves drive heterozygosity, which they do not for reasons given above, heterozygosity will be largely uncoupled from the component of D that reflects introgression.

It is noticeable that the relationship between heterozygosity and D is not restricted to African – non-African comparisons, but is instead seen across all comparisons, including those between populations within each major geographic region. This ubiquity extends even into Africa, where introgression is believed to be modest (Chen *et al*. 2020) or, in some populations, absent (Mallick *et al*. 2016). Across all these scenarios, informative sites lying in genomic regions where heterozygosity is higher one population tend to yield D values indicating introgression into the other population. Interestingly, this is also the pattern seen in simulated data with no introgression. In simulations, the mechanism is clear and involves ancestral human sequences tending to be commoner than derived sequences, such that homozygous regions are enriched for sequences with incomplete lineage sorting. When introgression is added to the simulations, this pattern reverses in comparisons between non-African populations or is dominated by a general increase in D in African – non-African comparisons. This contrasts strongly with empirical data, where non-zero D appears to be generated, not by a change in the size of D in any given region of the plot, but more by an asymmetrical distribution of the data across the diagonal, reflecting higher heterozygosity in one population compared with the other. In other words, real data follow an accentuated, asymmetrical version of the pattern seen where introgression is lacking.

The ubiquity of the relationship between flanking heterozygosity and D extends both to using the Denisovan as the archaic test group and to the use of only indels to measure heterozygosity, though the trend here is somewhat diluted. The influence of indels is interesting because small indels are known to be locally mutagenic (Tian *et al*. 2008), making the current results consistent with a model based on heterozygote instability. As with heterozygosity based on SNPs, the trend for indels is most strongly and consistently seen where data density is highest, i.e. at low heterozygosity values and closer to rather than further from the diagonal. Although weaker, the fact that SNPs and indels exhibit related patterns is important because indels do not contribute in any way to D, so provide an entirely independent signal.

The current results thus suggest that positive D is dominated by sites that are homozygous in non-Africans, these tending to be closer to the ancestral state and hence to related lineages like Neanderthals. In this sense, the current results offer the flip side of a previous analysis that showed how the signal captured by D was dominated by sites that are heterozygous in Africa acting to increase divergence from the human common ancestor, and hence related lineages (Amos 2020). There appear only two mechanisms capable of generating non-zero D: introgression and variation in mutation rate between populations, the latter having previously been assumed not to exist (Green *et al*. 2010; Durand *et al*. 2011; Patterson *et al*. 2012). If the link to heterozygosity can be shown not to be due to introgression then, by elimination, variation in mutation rate must be responsible. In this sense, the observed pattern is consistent with a model where loss of heterozygosity out of Africa caused a reduction in mutation rate (Amos 2013) which in turn reduced the rate of divergence from the ancestral state. Across the genome, natural selection and the stochasticity of drift have resulted in considerable variation in the amount of heterozygosity lost out of Africa and, in a few cases, heterozygosity actually increased. The amount of heterozygosity lost predicts measures of mutations rate (Amos 2013).

A further subtle but perhaps telling observation is the test for an effect of proximity. In simulated data, an increase in distance between where heterozygosity is measured and the focal base causes a general trend towards the genome-wide mean, as expected if D and heterozygosity are less and less coupled. In real data, a different pattern is observed, with the relationship with heterozygosity declining over a distance of only a handful of kilobases, an effect not seen in simulated data. In addition, the effect of heterozygosity is clearly *relative*, in the sense that the flip from positive to negative D occurs at the diagonal of equality. Since any given base in, say, Europe cannot ‘know’ whether the homologous base in Africa lies in a higher or lower heterozygosity context, there must be a direct link between heterozygosity and the generation of ABBAs and BABAs.

It could be argued that D is no longer the measure of choice for inferring introgression and that more sophisticated methods such as Conditional Random Field (CRF) (Mallick *et al*. 2016) should be preferred. However, all methods act on the same signal, the extent to which sequences in one population share more derived bases with an archaic relative to the same sequences in a second population. The D statistic is simple and has a strong theoretical basis (Durand *et al*. 2011; Martin *et al*. 2015). If introgression has occurred and left an appreciable legacy this should generate a signal that will be captured by D. The process of converting D into estimates of legacy size is less straightforward because it is strongly influenced by poorly known parameters such as population sizes and the times at which species / populations split. CRF attempts to use a simple model of sequence evolution to estimate the probability that any given base is of Neanderthal origin, and then integrates across the genome. While this method has much potential, its output can only be as accurate as the underpinning model of evolution used. Unfortunately, recent studies reveal a host of unexpected elements: mutations often occur in clusters of up to eight substitutions on the same strand (Besenbacher *et al*. 2016), recombination rate and mutation rate are correlated (Arbel-Eden and Simchen 2019; Halldorsson *et al*. 2019; Rousselle *et al*. 2019), mutation rates and spectra vary between populations(Amos 2013; Harris 2015), recombination hotspots show extreme temporal transience (Ptak *et al*. 2005). In addition, while it has previously been assumed that back-mutations can be ignored, in reality they appear common, with an estimated half a million sites across the autosomal genome (Amos 2020). These elements cannot yet be included in CRF calculations because their rates and any inter-dependencies have yet to be determined.

A model where heterozygosity acts to modulate mutation rate which in turn drives variation in the level of divergence from the ancestral state has a number of interesting implications. Most directly, it negates the need to invoke selection to explain why haploid genomic components like mitochondrial DNA and the Y chromosome show no evidence of introgression, while the semi-haploid X-chromosome shows much reduced evidence of introgression. Another observation that is consistent with this model is the link to immune related genes and other genes under selection. Natural selection has exerted a potent influence on the extent to which loss of heterozygosity out of Africa is either accelerated or reduced, particularly at immune-related genes. Consequently, we can expect to see unusually large (or small) D values in the vicinity of immune-related of other selected genes, just as observed.

In conclusion, heterozygosity measured within a kilobase of putative introgressed bases is found to be a potent and ubiquitous predictor of D, even in populations where introgression is thought absent. This relationship is not driven by introgressed fragments themselves, but instead seems to reflect an exaggerated form of the general tendency for ancestral sequences to be at high frequency and hence homozygous. I suggest a new model in which heterozygosity itself acts to modulate mutation rate which then in turn feeds back on this tendency to drive variation in D. Among other things, this new model can explain the pattern of rising D west to east across Eurasia (Wall *et al*.2013) which mirrors the decline in heterozygosity, the absence of introgressed haploid and semi-haploid regions, higher D values around genes under selection (Mendez *et al*. 2012; Quintana-Murci 2019) and why introgression is inferred almost wherever it is looked for (Kuhlwilm *et al*. 2019).

## Methods

### Data

Data were downloaded from Phase 3 of the 1000 genomes project (1000 Genomes Project Consortium 2010) as composite vcf files (available from ftp://ftp.1000genomes.ebi.ac.uk/vol1/ftp/release/20130502/). These comprise low coverage genome sequences for 2504 individuals drawn from 26 modern human populations spread across five geographic regions. Individual chromosome vcf files for the Altai Neanderthal were downloaded from http://cdna.eva.mpg.de/neandertal/altai/AltaiNeandertal/VCF/. Vcf files for the Denisovan genome were downloaded from http://cdna.eva.mpg.de/denisova/VCF/human/. I focused only on homozygote archaic bases, accepting only those with 10 or more reads, fewer than 250 reads and where >80% were of one particular base. This approach sacrifices modest numbers of (usually uninformative) heterozygous sites but benefits from avoiding ambiguities caused by coercing low counts into genotypes.

### Data Analysis

All analyses of the 1000g data were conducted using custom scripts written in C++ (ESM1). Since the 1000 genomes data are low coverage and include much imputation, population allele frequencies are determined with greater reliability than individual genotypes. Consequently, D statistics were calculated probabilistically assuming random assortment. D values were only calculated where the full autosomal genome yields a total of two or more counts for ABBA+BABA combined. To improve visualisation of the many near-zero values, D values are presented as the signed root of the raw value. Heterozygosity was calculated using two 1Kb windows placed symmetrically either side of but excluding each informative base that contributes to D. Note, each informative base is considered in isolation, in the sense that other informative bases in the flanking regions are used to calculate heterozygosity. Heterozygosity is calculated as the sum of the heterozygosities of all polymorphic sites within the two windows and is calculated separately for SNPs and indels. SNP heterozygosities are multiplied by four and then the integer value used to define 30 bins. Indel heterozygosities are smaller and were multiplied by eight before converting to 30 bins.

### Simulations

Simulated data were generated using the coalescent program ms (Hudson 2002). The base model was coded:

./ms 402 5000 -t 200 -I 6 1 1 100 100 100 100 -r 100 25000 -es 0.06 5 0.965 -ej 0.0601 7 2 - ej 0.015 6 5 -ej 0.03 4 3 -en 0.0685 5 0.007 -ej 0.07 5 3 -ej 0.5 3 2 -ej 6 2 1

I assume a haploid population size of 10,000, a mutation rate of 10^−8^ and set theta to 200, such that each of 5,000 recombining fragments is 500Kb long. The hominin-chimpanzee split is taken to be 6,000,000 years ago, the Neanderthal-human split is here 500,000 years ago (generation length = 25 years. The human population splits into African and non-African lineages 70,000 years ago, the non-African lineage immediately experiencing a bottleneck that removes about 25% of heterozygosity.

The African and non-African lineages each split into two at 30,000 and 15,000 years ago respectively. Introgression occurs at 60,000 years ago with a 3.5% injection into the non-African lineage.

## Acknowledgements

I thank Simon Martin, Rob Foley and Marta Lahr for many useful discussions.

